# Spatial specialization of antioxidant defenses dictates predation success in *Myxococcus xanthus*

**DOI:** 10.64898/2026.07.07.737134

**Authors:** Duo-hong Sheng, Xuan-qi Zhang, Xin-yao Yan, Li Zhuo, Yue-zhong Li

**Author notes:** **Corresponding author:** Duo-hong Sheng; Yue-zhong Li.

## Abstract

Redox-based chemical warfare is a primary driver of microbial community assembly. Here, we show that the predatory bacterium *Myxococcus xanthus* employs a spatial division of labor between two inducible monofunctional catalases, mxKatB and mxKatE, to overcome prey-derived hydrogen peroxide (H₂O₂). Quantitative transcript analysis revealed distinct regulatory specificities: mxkatB was the dominant transcriptional responder to exogenous H₂O₂, whereas mxkatE was preferentially induced by UV irradiation. Biochemical analyses demonstrated strict compartmentalization of enzymatic activity. mxKatE functioned intracellularly, consistent with a role in mitigating endogenous genotoxic stress. In contrast, mxKatB, which harbors an N-terminal Sec-dependent signal peptide, was exclusively localized to the extracellular milieu. Targeted gene deletions corroborated these non-redundant physiological roles. ΔkatE mutant exhibited severe growth defects and heightened sensitivity to UV and H₂O₂ yet retained full predation proficiency. Conversely, ΔkatB mutant displayed unaltered vegetative fitness but were severely impaired in prey lysis due to oxidative inactivation of secreted bacteriolytic enzymes. Failure of cross-complementation confirmed that spatial localization, rather than catalytic capacity, dictates enzyme function. Our findings establish that *M. xanthus* deploys an extracellular catalase shield to protect its exoenzyme arsenal from prey-derived oxidants. This spatial specialization of antioxidant defenses represents a sophisticated strategy that directly determines the outcome of bacterial predation and shapes interspecies interactions within microbial communities.

**IMPORTANCE:** Predatory bacteria such as *M. xanthus* must withstand chemical defenses deployed by their prey. We show that *M. xanthus* uses a spatially specialized antioxidant system: an extracellular catalase (mxKatB) secreted to shield its lytic enzymes from prey-derived hydrogen peroxide, and an intracellular catalase (mxKatE) that handles endogenous oxidative stress. This division of labor reveals that bacterial antioxidant defenses can be compartmentalized to protect extracellular weaponry rather than the cell itself, adding a new dimension to how spatial organization of stress responses influences the outcome of microbial competition.

## INTRODUCTION

Microbes dominate virtually every ecosystem on Earth, engaging in relentless chemical warfare to secure limited resources and physical space (1, 2) To gain a competitive advantage, bacteria have evolved diverse antagonistic strategies, ranging from the secretion of diffusible toxins—such as antibiotics, bacteriocins, and hydrogen peroxide (H_2_O_2_)—to contact-dependent killing via molecular syringes like the type VI secretion system (T6SS) (3–6). A common consequence of these interspecies conflicts is the induction of reactive oxygen species (ROS) in target cells (7–9). Among these, H_2_O_2_ serves as a particularly potent weapon. Unlike the short-lived superoxide anion (O2·−) or the hydroxyl radical (·OH), which are constrained by charge repulsion and membrane impermeability (10, 11), H_2_O_2_ is relatively stable and diffusible. It readily traverses cell membranes to act as both an oxidative stressor and a signaling molecule (12, 13). Furthermore, in the presence of labile iron, H_2_O_2_ drives the Fenton reaction, generating highly reactive ·OH that indiscriminately damages DNA, lipids, and proteins (14–16). Consequently, H_2_O_2_ is widely deployed as a chemical defense across biological kingdoms, from the oxidative burst of immune cells and plants to the antimicrobial arsenal of lactic acid bacteria (17–20).

To survive in H_2_O_2_-rich environments, bacteria rely on catalases—enzymes that disproportionally decompose H_2_O_2_ into water and oxygen (2 H_2_O_2_→2H_2_O+O_2_). While catalases are phylogenetically diverse, comprising monofunctional heme catalases, catalase-peroxidases, and manganese catalases (21–23), the monofunctional heme catalases are ubiquitous in aerobic microbes (24). Phylogenetic analyses suggest that these enzymes underwent gene duplications, leading to three major clades: Clade 1 (small subunit, plant/bacterial), Clade 2 (large subunit, KatE-like), and Clade 3 (small subunit, broad distribution) (23, 25).

The predatory bacterium *Myxococcus xanthus* employs two such monofunctional catalases, mxKatB (Clade 1) and mxKatE (Clade 2) (25, 26). Previous studies suggested that mxKatE functions as a housekeeping enzyme, while mxKatB plays a complementary role in H_2_O_2_ degradation (26). However, the functional partitioning of these enzymes during the complex process of predation remains unclear. Given that prey bacteria actively secrete H_2_O_2_ to resist predation, we hypothesized that *M. xanthus* utilizes the spatial specialization of these two catalases to mitigate oxidative damage. Here, we demonstrate a strict division of labor: mxKatE operates intracellularly to ensure stress resilience and growth, whereas mxKatB is exported to the extracellular milieu to shield secreted bacteriolytic enzymes, thereby overcoming prey oxidative defenses.

## MATERIALS AND METHODS

### Bacterial strains, plasmids, and culture conditions

The bacterial strains and plasmids used in this study are listed in Tables S1 and S2. *Escherichia coli* and *Salmonella enterica* were cultured in Luria-Bertani (LB) medium at 37 °C. *M. xanthus* strains were grown in CYE (1% Bacto-casitone, 0.5% Yeast Extract, 0.1% MgSO4. 7H_2_O, pH 7.6) liquid medium with shaking (200 rpm) or on CYE agar plates at 30 °C. Predation and developmental assays were conducted on TPM agar plate (10 mM Tris-HCl, pH 7.6; 8 mM MgSO₄; 1 mM KH₂PO₄; 1.5% agar). Antibiotics were supplemented where appropriate at the following concentrations: kanamycin (40 µg/mL) and ampicillin (100 µg/mL).

### Growth, development, and stress resistance assays

For phenotypic analyses, *M. xanthus* strains were pre-cultured in CYE medium to an optical density at 600 nm (OD₆₀₀) of 0.5, harvested by centrifugation, and washed with 10 mM phosphate buffer (pH 7.0).

#### Growth and Development

Cells were adjusted to 0.02 OD₆₀₀ in CYE liquid medium and incubated for 96 h; growth was monitored by measuring OD₆₀₀ every 12 h. For fruiting body formation, 1.5 × 10⁹ cells were spotted onto TPM agar plates (14 cm diameter). Myxospores were isolated from mature fruiting bodies, heat-treated at 60 °C for 2 h to kill residual vegetative cells, and briefly sonicated. Germination assays were conducted in CYE plate at 30 ℃ for 5 days and then counted.

#### Stress Resistance

For UV exposure, cell suspensions in phosphate buffer were irradiated using a UV Crosslinker (Fisher Scientific) at doses ranging from 0 to 25 J/m². For oxidative stress, cells were exposed to 0–10 mM H₂O₂ in phosphate buffer for 20 min at room temperature. Then, the cells were re-suspended in fresh CYE medium and incubated at 30 °C for 0-4h. Survival rates were determined via soft agar colony formation assays. Briefly, serial dilutions were mixed with 0.6% CYE soft agar and overlaid onto CYE plates. Colonies were counted after incubation, and survival percentages were calculated relative to untreated controls.

### Genetic manipulation

Chromosomal DNA from *M. xanthus* and plasmid DNA from *E. coli* were extracted using standard kits (Tiangen). For heterologous expression, *mxkatB*, *Δsp_katB* (encoding a signal-peptide deletion variant), and *mxkatE* were amplified and cloned into the pET29b vector using the ClonExpress II One Step Cloning Kit (Vazyme).

Markerless deletion mutants (ΔKatB and ΔKatE) were constructed in *M. xanthus* DK1622 using the suicide vector pBJ113, which carries a kanamycin resistance cassette for positive selection and the *galK* gene for negative selection (27). Upstream and downstream homologous arms were ligated into pBJ113 and introduced into *M. xanthus* via electroporation (1.25 kV, 400 Ω and 1 mm cuvette). Double recombinants were selected on CYE plates containing 1% galactose. Complementation and overexpression constructs were derived from the pSWU19 plasmid (28) and introduced into strains via electroporation.

### RNA extraction and gene expression analysis

Total RNA was extracted using RNAiso Plus (Takara) and treated with gDNA Eraser to remove genomic DNA contamination. First-strand cDNA was synthesized using the PrimeScript™ RT Reagent Kit (Takara). Quantitative real-time PCR (qPCR) was performed on an ABI StepOnePlus system using SYBR Premix Ex Taq (Takara). Gene expression was normalized to *gapA* and calculated using the 2⁻^ΔΔCt^ method (29). Semi-quantitative PCR was performed using gene-specific primers.

For transcriptomics, RNA sequencing was conducted by Vazyme (Nanjing, China) with three biological replicates per condition (Table S3). Differentially expressed genes were annotated against the NR and KEGG databases.

### Protein expression, purification, and enzymatic assays

Recombinant proteins were expressed in *E. coli* BL21 (DE3) by induction with 0.1 mM isopropyl β-D-1-thiogalactopyranoside (IPTG) at 16 °C for 20 h. Cells were lysed by sonication in lysis buffer (25 mM Tris-HCl, pH 8.0; 200 mM NaCl; 5% glycerol), and soluble fractions were purified using Amylose resin (New England Biolabs) according to the manufacturer’s protocol.

To analyze native catalase activity, *M. xanthus* cells were separated from culture supernatants by centrifugation. Cell pellets were disrupted by sonication, and extracellular proteins in the culture supernatant were precipitated with 60% saturated ammonium sulfate. Protein concentrations were determined using a BCA assay (Vazyme). Native PAGE (7.5% gel) was performed at 4 °C, and catalase activity was visualized using a staining method adapted from Sun et al. (30). Quantitative catalase activity was measured using an Ammonium Molybdate Colorimetry Kit (Jiancheng Bioengineering Institute), monitoring the decomposition of H₂O₂ at 405 nm.

### Predation assays

Predator and prey cells were grown to mid-log phase, washed, and adjusted to 1.0 OD₆₀₀ in TPM buffer. Predation efficiency was evaluated using two assays: distance predation assays (DPAs) for qualitative analysis and overlapping predation assays (OPAs) for quantitative measurements (31–33).

### Structural modeling and docking

Three-dimensional structures of mxKatB, Δsp_KatB (signal peptide truncated), and mxKatE were predicted using AlphaFold-Server (https://alphafoldserver.com) and the SWISS-MODEL homology modeling pipeline (https://swissmodel.expasy.org/). Potential protein-protein interactions and structural connectivity were analyzed using the SmoothDock server (http://structure.pitt.edu/servers/smoothdock). Structural visualizations were generated using PyMOL.

### Data availability and biosafety

All raw sequencing data have been deposited in the NCBI Sequence Read Archive under BioProject accession PRJNA1474103. All source data underlying the figures are available in the supplementary material. All experiments complied with local biosafety regulations.

## RESULTS

### Transcriptomic profiling reveals induction of a secreted catalase during predation

On nutrient-free TPM agar, *M. xanthus* efficiently lysed *S. enterica* colonies within 72 h (Fig. 1A). To mimic oligotrophic natural environments, we added 50 mM glucose—a treatment that enhances prey biomass without disrupting *Myxococcus* development (34)—which markedly suppressed predation (Fig. 1A). This suggested that nutrient availability alters the chemical warfare dynamics between predator and prey.

**Figure 1.**
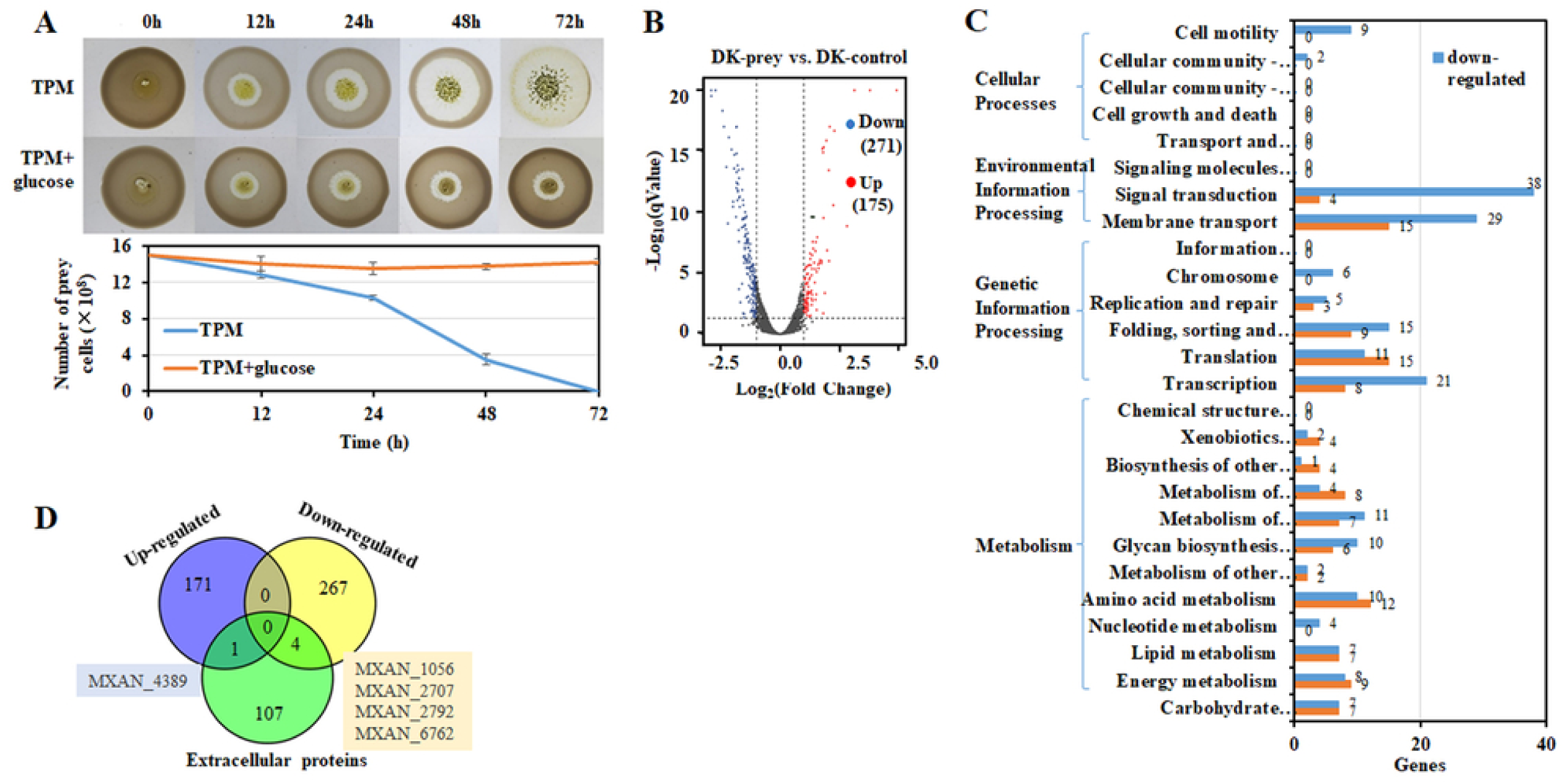
Transcriptomic profiling of *Myxococcus xanthus* during predation of Salmonella enterica. **(A)** Validation of predation by *M. xanthus* on *S. enterica. S. enterica* (1.5 × 10⁹ CFU) was spotted onto TPM agar plates, and *M. xanthus* (10⁶ CFU) was inoculated at the center. After incubation at 32 °C for 0–72 h, surviving prey cells were enumerated by plate counting (lower panel). Data are means ± SD (n = 3). **(B)** Volcano plot of differentially expressed genes (DEGs) identified by RNA-seq between DK-prey and DK-control. The x-axis shows log_₂_(fold change) (log₂FC), and the y-axis shows −log₁₀(adjusted P value). Genes with log₂FC ≥ 1 and adjusted P < 0.05 were considered significantly differentially expressed. Red points represent 175 significantly upregulated genes, blue points represent 271 significantly downregulated genes, and gray points represent genes not meeting the significance threshold. **(C)** KEGG pathway enrichment analysis of DEGs induced during predation. *M. xanthus* cDNA libraries were prepared after 2 h of co-culture with *S. enterica* in TPM medium—representing an optimal window for elucidating predation mechanisms (Pérez et al., 2022). Significantly enriched pathways (adjusted *P*< 0.05) are displayed. **(D)** Venn diagram illustrating the overlap between predation-induced DEGs and the previously reported extracellular proteome of *M. xanthus* (Evans et al., 2012).

To elucidate the molecular basis of predation, we compared the transcriptomes of *M. xanthus* under starvation versus during predation on *S. enterica*. Principal component analysis confirmed high reproducibility between biological replicates (Supplementary Fig. S1). Compared to starved controls, co-culture with prey elicited differential expression of 446 genes (175 up-regulated, 271 down-regulated) (Fig. 1B) (Table S4, S5). Down-regulated genes were enriched in motility and signal transduction, whereas up-regulated genes were associated with translation and xenobiotics metabolism (Fig. 1C). Notably, comparative analysis of our transcriptome against published secretomes datasets (35) identified only one extracellular protein significantly induced during predation: **MXAN_4389**, encoding a monofunctional catalase designated mxKatB (Fig. 1D).

### Prey-derived H₂O₂ acts as a chemical barrier against predation

Given the induction of a catalase during predation, we hypothesized that *S. enterica* deploys H₂O₂ as an oxidative defense. While neither organism produced detectable H₂O₂ in nutrient-free conditions, both *S. enterica* ST11 and *E. coli* 1655 secreted H₂O₂ in the presence of glucose (Fig. 2A). Crucially, H₂O₂ production by *S. enterica* was triggered by the proximity of *M. xanthus*, forming a distinct halo around the prey colony that intensified upon contact (Fig. 2B). Quantitative analysis revealed that direct contact induced H₂O₂ concentrations reaching up to 2 mM at the colony edge.

**Figure 2.**
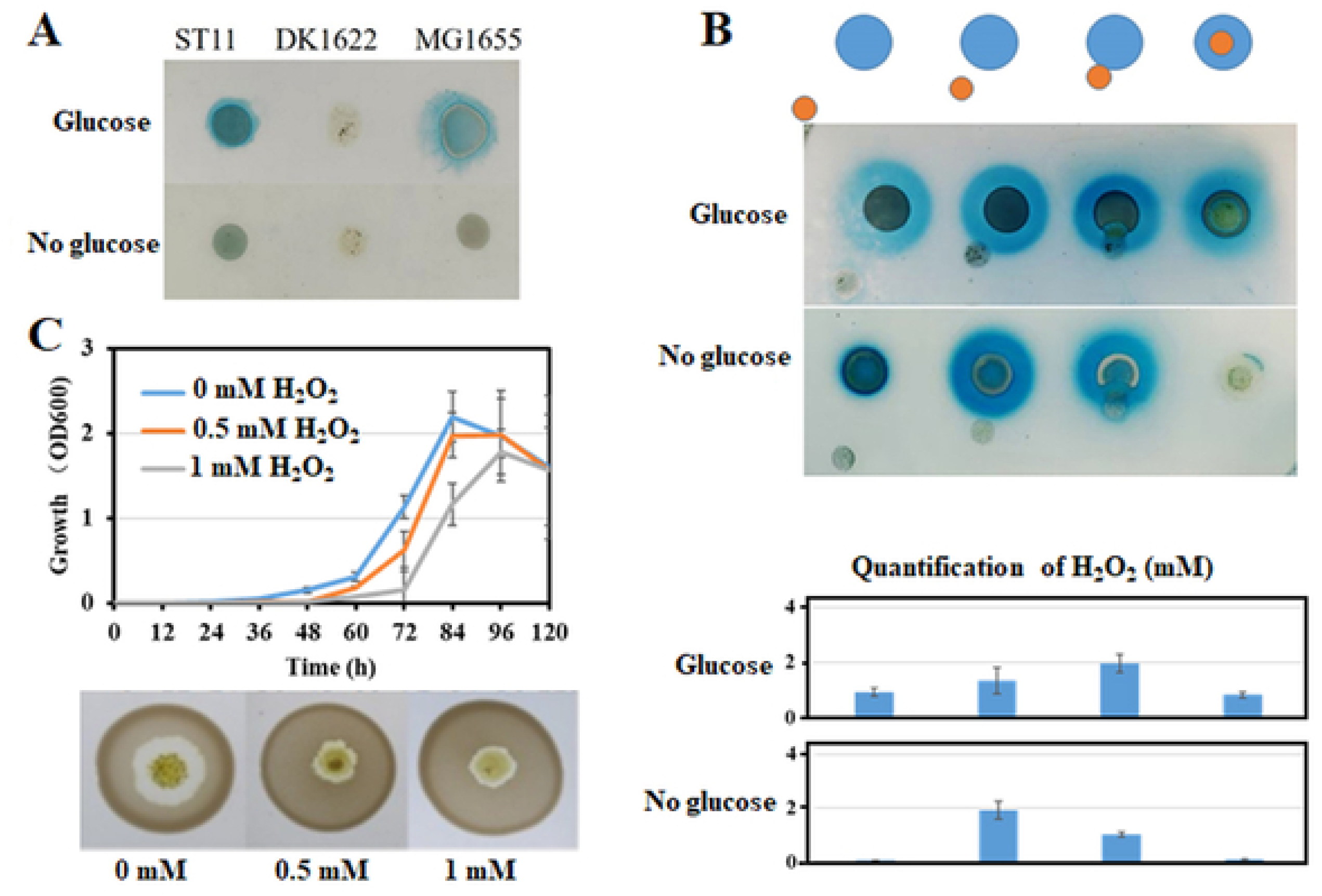
Extracellular H₂O₂ production and its impact on *M. xanthus*. **(A)** Extracellular H₂O₂ production by *S. enterica* (ST11), *M. xanthus* (DK1622), and *E. coli* (MG1655), detected by potassium ferricyanide–ferric chloride staining. **(B)** Predation-induced H₂O₂ production. *M. xanthus* and *S. enterica* were co-cultured at the indicated spatial configurations (1 cm, 0.5 cm, 0 cm, overlapping) and incubated at 30 °C for 48 h. Agar surrounding prey colonies was harvested, centrifuged at 12,000 × *g* for 10 min, and supernatants were used for H₂O₂ quantification. **(C)** Effects of gradient concentrations of H₂O₂ on *M. xanthus* growth and predation efficiency. Error bars denote SD from three independent experiments (*n*= 3).

Exogenous H₂O₂ severely compromised *M. xanthus* fitness; concentrations as low as 0.5 mM delayed growth and reduced predation efficiency, while 1 mM H₂O₂ completely abolished predatory activity (Fig. 2C). These results establish **H₂O₂ as a potent anti-predation weapon employed by *S. enterica*.**

### Distinct regulatory architectures of the two catalases

*M. xanthus* encodes two catalases: *mxkatB* and *mxkatE*. Promoter analysis revealed distinct regulatory logics (Fig. 3A). *MxkatB* harbors five putative binding sites for PexR, a transcriptional activator known to sense H₂O₂ (36). Conversely, *mxkatE* lacks PexR sites but contains a SoxR-binding motif, suggesting responsiveness to superoxide stress (37).

**Figure 3.**
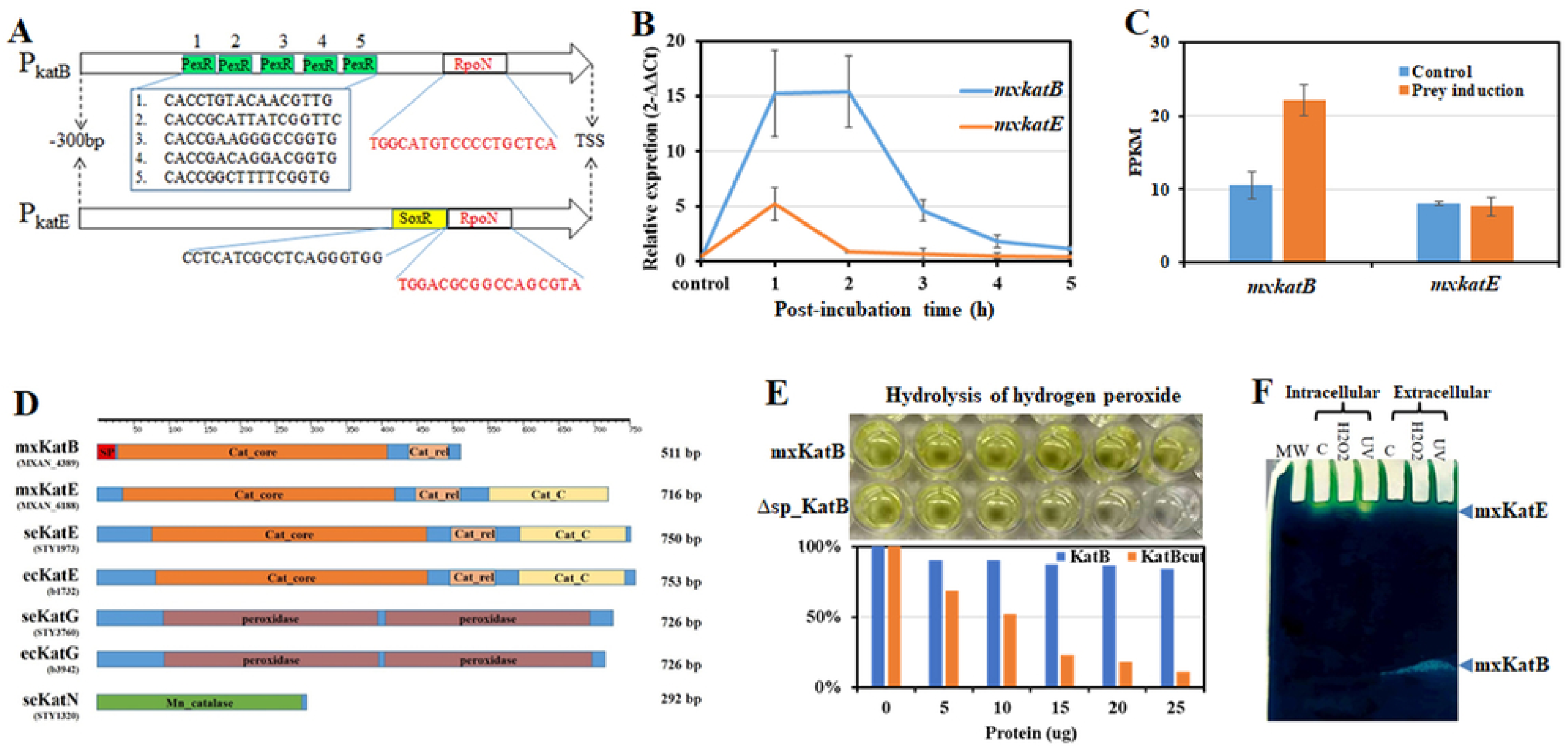
Transcriptional regulation and biochemical characterization of two catalases in *M. xanthus*. **(A)** Schematic of the functional promoter regions (−300 to 0 bp) of *mxkatB* and *mxkatE*. Predicted binding sites for the alternative sigma factor σ⁵⁴ (RpoN), PexR, and SoxR are indicated by red text, green shading, and yellow shading, respectively. **(B)** RT–qPCR analysis of *mxkatB* and *mxkatE* transcription in *M. xanthus* exposed to 2 mM H₂O₂. Relative mRNA levels were normalized to an internal reference gene. **(C)** Transcriptomic expression of mxKatB and mxKatE pre- and post-predation. Values are means ± SD (n= 3). **(D)** Domain architecture comparison of catalases from *S. enterica*, *M. xanthus*, and *E. coli*. Predicted signal peptides are shown in red. **(E)** Catalytic activity of recombinant mxKatB and Δsp-KatB towards H₂O₂. Upper panel: residual H₂O₂ staining; lower panel: quantitative measurement of residual H₂O₂. **(F)** *In-gel* catalase activity staining of intracellular and extracellular fractions from *M. xanthus*. Log-phase cells were treated with 2 mM H₂O₂ or 10 J / m² UV, followed by incubation at 30 °C for 1 h. Proteins were extracted, quantified, and separated by native PAGE.

Consistent with this, exposure to 2 mM H₂O₂ induced *mxkatB* expression over 15-fold, with detectable induction occurring within 1 h (Fig. 3B). In contrast, *mxkatE* was only marginally induced (<2-fold) (Fig. 3B). Transcriptomic data further corroborate that mxKatB serves as the primary enzyme responsible for the rapid response to H₂O₂ (Fig. 3C).

### Spatial segregation of catalase activity

Structural analysis revealed that mxKatB lacks the C-terminal domain present in mxKatE but possesses a unique N-terminal extension predicted to be a Sec-dependent signal peptide (SP) (Fig. 3D, Supplementary Fig. S3-S9). Heterologous expression in *E. coli* showed that full-length mxKatB exhibited weak enzymatic activity, whereas a truncated variant lacking the SP (Δsp_KatB) displayed a ∼7-fold increase in H₂O₂ degradation (Fig. 3E). This suggests the SP maintains the enzyme in a translocation-competent, partially inactive state (38, 39).

Subcellular fractionation confirmed this spatial division: mxKatE activity was strictly intracellular, while mxKatB activity was exclusively detected in the extracellular fraction (Fig.3F). H₂O₂ exposure specifically upregulated extracellular mxKatB activity without affecting intracellular mxKatE.

Phylogenetic analysis of 497 KatB homologs across 9,959 bacterial genomes (GenomeNet database, release June 2026) revealed a striking pattern: Gram-positive bacteria (Bacillota) possess KatB without SPs, whereas Gram-negative bacteria (including Myxococcota) typically encode KatB with SPs (Supplementary Fig. S11A–B). Notably, predatory genera within Myxococcota exhibited the highest prevalence of SP-containing KatB (>42.5%), primarily in predatory genera such as *Myxococcus* and *Corallococcus* (Supplementary Fig. S11C), implying a specialized role in interspecies interactions.

### Functional specialization in growth, stress resistance, and predation

We constructed deletion mutants to dissect their physiological roles (Supplementary Fig. S12). Deletion of *mxkatE* severely impaired growth, extending the lag phase and reducing biomass (Fig. 4A), confirming its role as a housekeeping enzyme. In contrast, ΔkatB grew similarly to the wild type. Neither mutation affected fruiting body development or sporulation (Fig. 4B).

**Figure 4.**
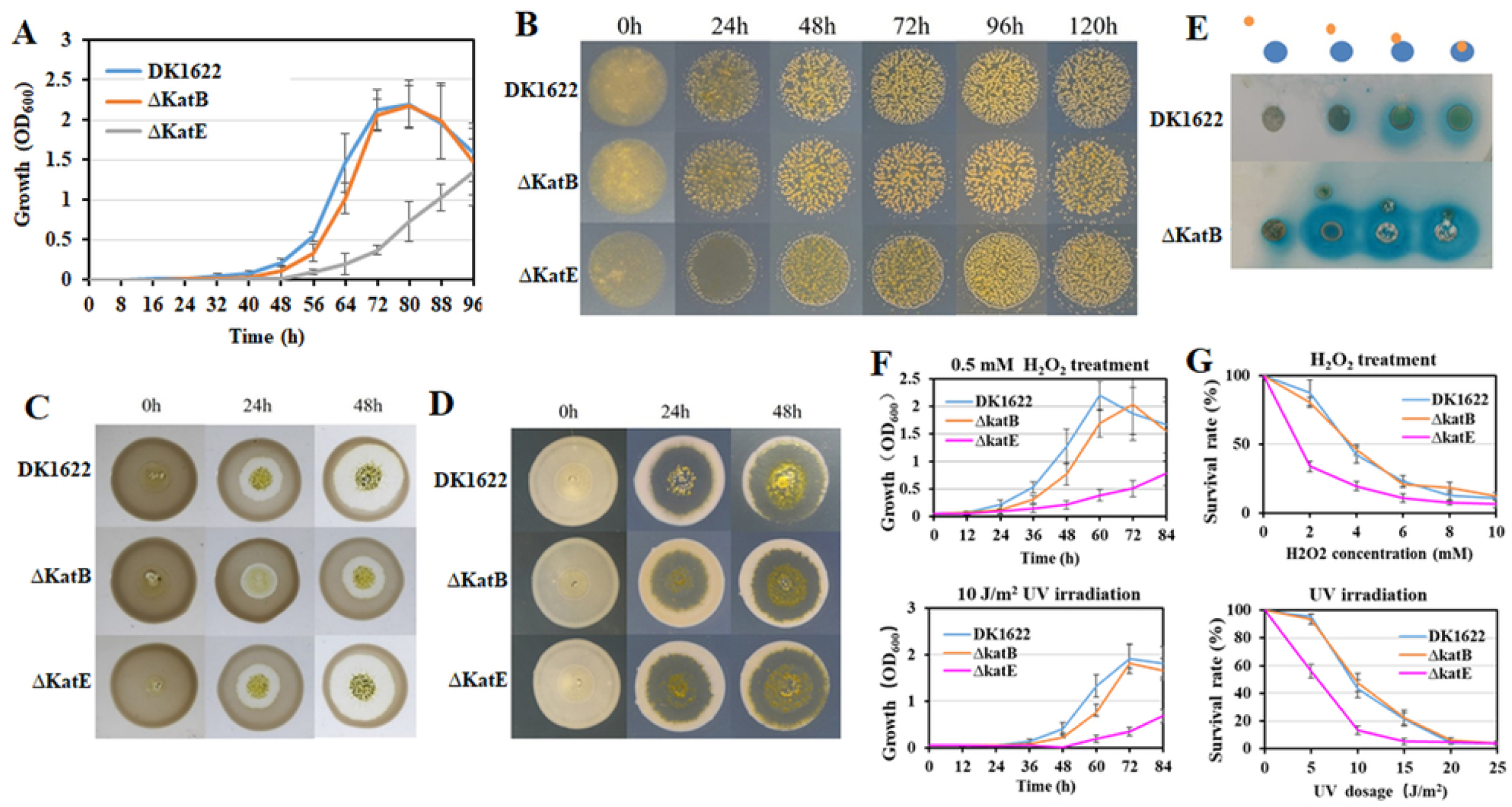
Phenotypic characterization of catalase-deficient mutants in *M. xanthus*. **(A)** Growth curves of wild-type DK1622 and catalase mutants (ΔKatB and ΔKatE). Data are means ± SD (*n*= 6). **(B)** Developmental phenotypes of wild-type and mutant strains following 5 days of incubation at 32 °C. Images are representative of at least three biological replicates per strain. **(C)** Predation of live *S. enterica* by *M. xanthus*. Prey cell lysis was monitored at indicated time points under 15× magnification. **(D)** Predation on heat-killed prey cells. *Salmonella* was inactivated at 65 °C for 30 min, washed with TPM buffer, and adjusted to OD₆₀₀ = 1.0 prior to use. **(E)** Residual H₂O₂ staining during predation at varying spatial distances. DK1622 and ΔKatB were inoculated at distances of 1 cm, 0.5 cm, 0 cm, or in an overlapping configuration with prey cells. **(F)** Growth of wild-type and catalase mutants in CYE medium supplemented with 0.5 mM H₂O₂ or under chronic UV exposure (5 J/m² every 12 h). OD₆₀₀ was recorded every 12 h. **(G)** Sensitivity of mutants and wild-type to acute H₂O₂ and UV stress, assessed as described in Materials and Methods. Each data point represents the mean ± SD (*n*= 5).

However, predation assays revealed a stark functional divide. While ΔKatE exhibited wild-type predation efficiency, ΔkatB showed a significant reduction in prey lysis (Fig. 4C). This defect was specific to live prey; both mutants predated equally well on heat-killed *S. enterica* (Fig. 4D), indicating that mxKatB neutralizes a labile, prey-derived defensive factor—namely, H₂O₂ (Fig. 4E).

Given the established roles of catalases in antioxidant defense and UV resistance (40–42), we evaluated the physiological contributions of the two enzymes under stress. Under chronic exposure to H₂O₂ or UV irradiation, the ΔkatB exhibited a moderate growth defect, whereas the ΔkatE displayed a severe proliferation impairment (Fig. 4F). Acute stress assays revealed a distinct division of labor: while ΔkatB survival rates were indistinguishable from the wild type, ΔkatE viability was markedly compromised, decreasing to approximately 50% under 2 mM H₂O₂ and 59% under 5 J/m² UV (Fig. 4G). Thus, **mxKatE ensures intracellular self-protection, while mxKatB safeguards the extracellular predatory apparatus.**

### Lack of cross-complementation confirms non-redundancy

To further delineate the functional specificity of mxKatB and mxKatE, we performed cross-complementation assays using their native promoters (∼300 bp upstream). Constructs encoding full-length *mxkatB*, a signal-peptide truncation (*Δsp_katB*), and *mxkatE* were integrated into the chromosome via the pSWU19 vector (Fig. 5A-B). Quantitative PCR confirmed robust transcription of all transgenes (Fig. 5C).

**Figure 5.**
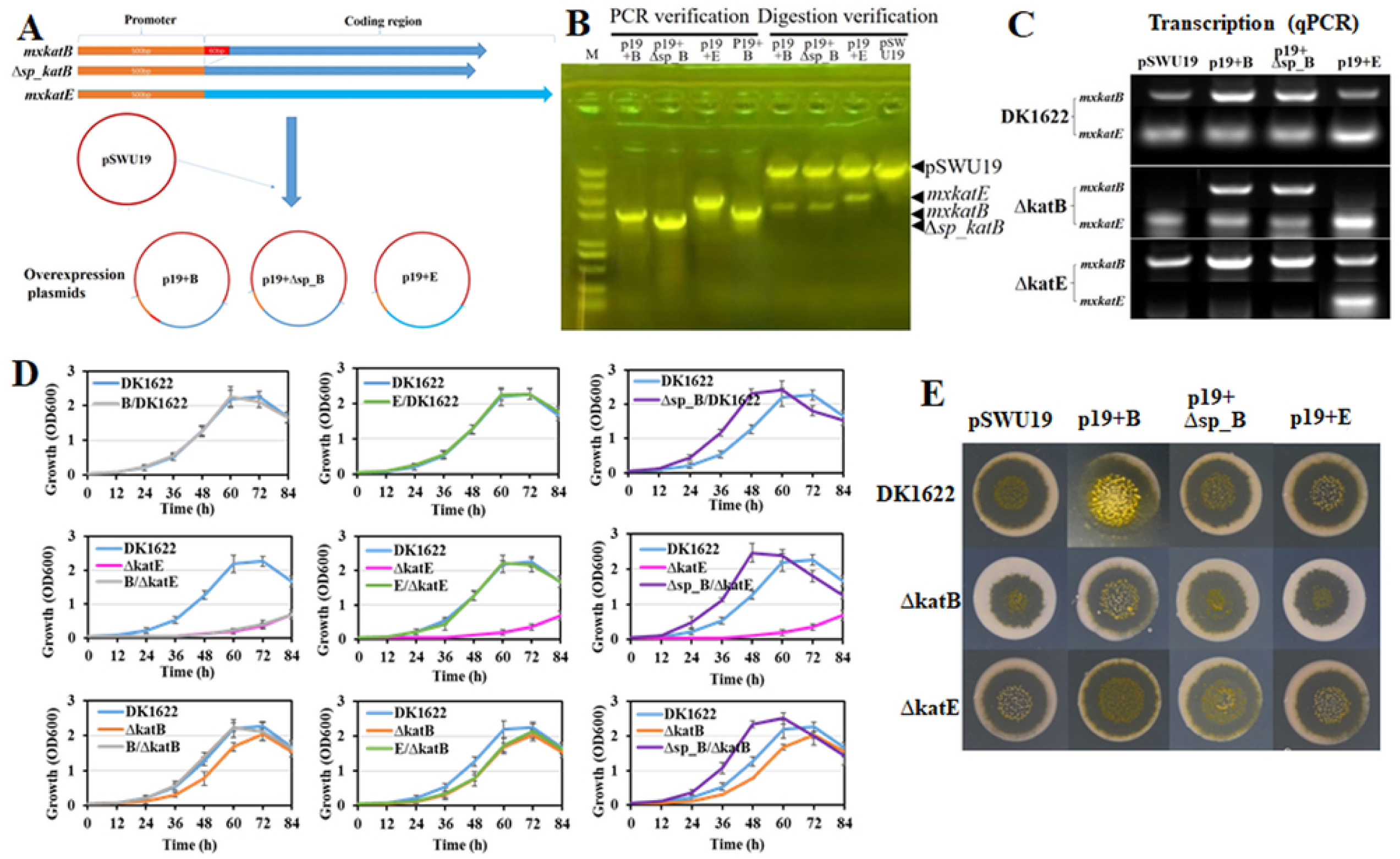
Genetic complementation and cross-compensation analysis of catalases. **(A)** Schematic of integrative expression plasmid construction. The *mxkatB* gene with its native 300-bp upstream region (primers Pb_F/katB_R) or the *mxkatE* gene with its 300-bp upstream region (primers Pe_F/katE_R) was cloned into the integration vector pSWU19, generating plasmids P19+B and P19+E, respectively. A truncated *katB* construct (Δ*sp_katB*) was generated by fusing the Δ*sp_katB* coding sequence (primers Pb_F/Pb_R) with its 300-bp upstream region (primers Bcut_F/katB_R) and inserted into pSWU19, yielding plasmid P19+Δsp_B. **(B)** Verification of recombinant plasmids by PCR amplification and restriction digestion analysis. **(C)** Semiquantitative RT–PCR analysis of *mxkatB* and *mxkatE* transcription in wild-type DK1622 and catalase mutants (ΔKatB, ΔKatE) harbouring the different expression plasmids. **(D)** Growth curves of complemented and overexpression strains. Data are means ± SD (*n*= 5). **(E)** Predation performance of complemented and overexpression strains against *S. enterica*.

In a wild-type background, overexpression of *mxkatB* or *mxkatE* had negligible effects on growth, whereas *Δsp_katB* significantly accelerated growth kinetics and increased biomass (Fig. 5D). Complementation assays revealed a strict functional segregation: p19+B and p19+E specifically rescued the growth defects of the ΔKatB and ΔKatE mutants, respectively, while cross-complementation failed. This reinforces that the two catalases operate via distinct mechanisms dictated by their cellular localization. Notably, expression of *Δsp_katB* conferred a growth advantage even in the mutant backgrounds, suggesting that releasing the enzyme from its translocation-incompetent state enhances its protective capacity.

Predation assays corroborated this specialization. p19+B fully restored the predation deficiency of ΔKatB and even enhanced wild-type predation efficiency (Fig. 5E). In contrast, p19+*Δ*sp_B failed to complement the predation defect, demonstrating that the **extracellular localization of mxKatB is indispensable for its role in overcoming prey defenses.**

### mxKatB protects extracellular lytic enzymes from oxidative inactivation

To determine how mxKatB enhances predation, we assayed extracellular lytic activity. Total extracellular proteins from *M. xanthus* lysed *S. enterica* cells in vitro, whereas protein-depleted supernatant did not (Fig. 6A), confirming that secreted lytic enzymes mediate killing.

**Figure 6.**
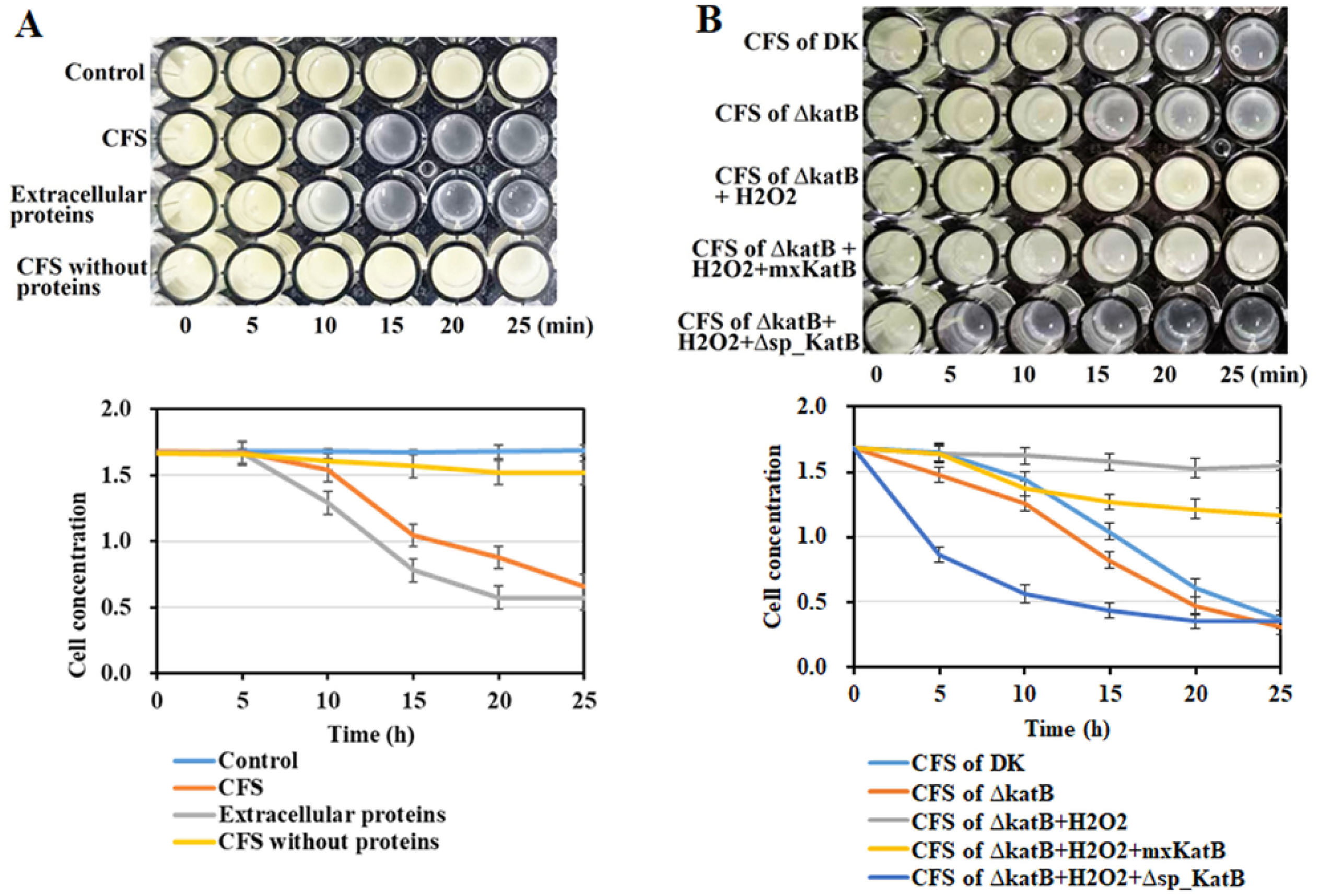
Contribution of extracellular catalase to prey cell lysis. Upper panels: representative images of prey cells at indicated time points (0–25 min). Lower panels: quantitative analysis of prey cell density. Error bars denote SD from three independent measurements. **(A)** Cell-free supernatant (CFS) of *M. xanthus* mediates prey lysis. **(B)** The signal peptide–truncated catalase Δsp-KatB preserves the lytic activity of CFS against H₂O₂-mediated inhibition.

While supernatants from wild-type and ΔKatB strains showed similar baseline lytic activity, the addition of 2 mM H₂O₂ selectively inhibited the lytic activity of ΔKatB supernatant (Fig. 6B). This inhibition was rescued by supplementing the reaction with the highly active *Δ*sp_KatB protein. Therefore, **extracellular mxKatB (Δsp_KatB) functions as a pericellular shield, protecting secreted bacteriolytic enzymes from oxidative inactivation by prey-derived H₂O₂.**

## DISCUSSION

Bacteria deploy an extensive chemical arsenal to compete for limited resources. Here, we define a spatial division of labor between two catalases that enables the predatory bacterium *M. xanthus* to overcome prey-derived oxidative defenses. We show that mxKatE functions as a conventional intracellular enzyme, mitigating endogenous and environmentally imposed oxidative stress to maintain cellular viability. In contrast, mxKatB is exported to the extracellular milieu, where it neutralizes prey-secreted H₂O₂ and preserves the activity of secreted bacteriolytic enzymes. This compartmentalization explains how a single predator manages intracellular redox homeostasis while simultaneously conducting chemical warfare at the cell surface.

Our data challenge the prevailing view that bacterial catalases are largely redundant or strictly intracellular. Although many bacteria encode multiple catalases, their functional specialization is often attributed to differences in substrate affinity or stress regime rather than subcellular localization (21–24). In *M. xanthus*, mxKatB and mxKatE are non-redundant: mxKatE is essential for growth and survival under oxidative stress, while mxKatB is dispensable for vegetative fitness but critical during predation. The strict failure of cross-complementation—particularly the inability of intracellular mxKatE to rescue the predation defect of ΔkatB—demonstrates that localization, not catalytic capacity, dictates physiological role.

The export of mxKatB represents a strategic investment. By maintaining catalase activity outside the cytoplasm, *M. xanthus* protects its extracellular lytic enzymes from inactivation by prey-derived H₂O₂. This mechanism parallels eukaryotic immune strategies, such as the secretion of catalase by fungal pathogens to disarm host oxidative bursts (43), but is notable in a bacterium that lacks a classical secretory system for oxidative defense. The presence of a Sec-dependent signal peptide in mxKatB appears to maintain the enzyme in a partially inactive, translocation-competent state until exported—a regulatory feature that may prevent premature H₂O₂ consumption before contact with prey.

Phylogenetic analysis supports an evolutionary link between catalase export and predation. Signal-peptide-bearing KatB orthologs are rare among bacteria but enriched in predatory Myxococcota, particularly within *Myxococcus* and *Corallococcus*. This distribution suggests that extracellular KatB arose as an adaptation to interspecies conflict rather than as a general stress-response enzyme. In this context, mxKatB can be viewed as a “cooperative enzyme,” functioning not to protect the producing cell directly but to preserve public goods—the secreted lytic enzymes—that benefit the local predator population.

These findings broaden our understanding of bacterial competition. Most studies of interbacterial antagonism focus on weapons—such as antibiotics, T6SS effectors, or ROS—rather than the protective systems that enable sustained attack. Our work highlights extracellular antioxidant defense as a critical component of microbial offensive capability. Because prey bacteria actively induce H₂O₂ upon predator contact, the ability to degrade extracellular ROS is likely a widespread, yet underappreciated, determinant of predation success.

Several questions remain. It is unclear whether mxKatB is tethered to the outer membrane or freely diffuses in the extracellular matrix, and whether other predators employ similar strategies. Additionally, the regulatory circuit linking prey detection to PexR-mediated mxkatB induction warrants further dissection. More broadly, our results suggest that the efficacy of predatory bacteria in natural ecosystems—such as soil or rhizosphere—may depend not only on their weaponry but also on their capacity to neutralize environmental and prey-derived oxidative stress.

In summary, we demonstrate that *M. xanthus* partitions two inducible catalases across cellular compartments to achieve both self-protection and offensive efficacy. This spatial specialization underscores the sophistication of bacterial chemical warfare and provides a framework for understanding how microbial predators persist in competitive, oxidatively stressful niches.

## Data availability statement

All relevant data are within the paper and its supporting information files. The transcriptome sequencing data are available in the SRA database under project accession PRJNA1474103.

## Author contributions

DHS and YZL designed experiments; DHS, XQZ, XYY, and LZ performed the experiments; DHS and YZL wrote the paper. All authors helped revise the manuscript.

## ACKNOWLEDGEMENTS

We thank the Life and Environment Research Shared Instrumentation Platform at Shandong University for instrumental and technical support. We also thank Y. Zhang for technical assistance with transcriptome analysis, and YR. Wang for English editing of the manuscript.

